# Unbiased interrogation of memory B cells from convalescent COVID-19 patients reveals a broad antiviral humoral response targeting SARS-CoV-2 antigens beyond the spike protein

**DOI:** 10.1101/2021.01.27.428534

**Authors:** Jillian M. DiMuzio, Baron C. Heimbach, Raymond J. Howanski, John P. Dowling, Nirja B. Patel, Noeleya Henriquez, Chris Nicolescu, Mitchell Nath, Antonio Polley, Jamie L. Bingaman, Todd Smith, Benjamin C. Harman, Matthew K. Robinson, Michael J. Morin, Pavel A. Nikitin

**Affiliations:** Immunome, Inc., Exton, PA, U.S.A.

## Abstract

Patients who recover from SARS-CoV-2 infections produce antibodies and antigen-specific T cells against multiple viral proteins. Here, an unbiased interrogation of the anti-viral memory B cell repertoire of convalescent patients has been performed by generating large, stable hybridoma libraries and screening thousands of monoclonal antibodies to identify specific, high-affinity immunoglobulins (Igs) directed at distinct viral components. As expected, a significant number of antibodies were directed at the Spike (S) protein, a majority of which recognized the full-length protein. These full-length Spike specific antibodies included a group of somatically hypermutated IgMs. Further, all but one of the six COVID-19 convalescent patients produced class-switched antibodies to a soluble form of the receptor-binding domain (RBD) of S protein. Functional properties of anti-Spike antibodies were confirmed in a pseudovirus neutralization assay. Importantly, more than half of all of the antibodies generated were directed at non-S viral proteins, including structural nucleocapsid (N) and membrane (M) proteins, as well as auxiliary open reading frame-encoded (ORF) proteins. The antibodies were generally characterized as having variable levels of somatic hypermutations (SHM) in all Ig classes and sub-types, and a diversity of V_L_ and V_H_ gene usage. These findings demonstrated that an unbiased, function-based approach towards interrogating the COVID-19 patient memory B cell response may have distinct advantages relative to genomics-based approaches when identifying highly effective anti-viral antibodies directed at SARS-CoV-2.

## INTRODUCTION

As the one-year mark approaches since its emergence in Wuhan, China in late 2019 [1], the prolonged spread of the severe acute respiratory syndrome coronavirus 2 (SARS-CoV-2) virus has resulted in one of the most devastating global health challenges of the last century [2–4]. With greater than 45 million confirmed cases and nearly 1.2 million deaths world-wide (WHO, as of November 2020 [5]), the virus continues to pose an extraordinary challenge to the scientific community, consequently becoming an unprecedented socio-economic disaster and burdening healthcare systems around the world. Infection with SARS-CoV-2 results in a myriad pathologies [6] collectively referred to as COVID-19 [3]. While a majority of individuals who become infected with the virus are capable of generating a productive anti-viral response, for many their anti-viral humoral response will not be sufficient to shield them from a potentially deadly infection. Therefore, as the global community braces for the next spike in infection and mortality rates, the urgency to develop effective therapeutics recapitulating the productive anti-viral response to combat the swelling health crisis has never been greater.

SARS-CoV-2 genomic RNA contains a large viral replicase gene, genes encoding non-structural proteins at its 5’ end, and a region encoding four major structural and multiple accessory proteins at the 3’ end. Structural proteins include Spike or Surface glycoprotein (S), Membrane protein (M), Envelope protein (E) and Nucleocapsid protein (N) [7]. The membrane surface glycoprotein S consists of two subunits, S1 and S2, that mediate viral binding to the host receptor ACE2 and fusion with the host cell membrane, respectively. The S1 subunit contains the receptor binding domain (RBD) that directly interacts with ACE2 and is a target of multiple neutralizing antibodies currently in clinical trials [8,9].

Genetic analyses of immune effector cells from convalescent patients who had effectively cleared the SARS-CoV-2 virus revealed that these individuals often had robust T and B cell responses to multiple other viral antigens beyond S protein [10–14], suggesting that the recognition of multiple antigens beyond the S protein may be important for viral clearance and the efficient resolution of infection. Included among the targets of the adaptive immune response were N and M proteins, as well as proteins encoded by the viral open reading frame (ORF). As such, a multi-targeted mixture of high-affinity anti-SARS-CoV-2-specific antibodies, more reflective of the broad humoral response seen in high-titer, mild to moderate COVID-19 convalescent patients, may be a more effective therapeutic strategy than using S-specific antibodies alone.

This report describes studies to elucidate the memory B cell antibody response in convalescent patients, using a method that enables the generation of large, stable hybridoma libraries from primary human B cells. This approach was previously used to identify a panel of monoclonal antibodies from convalescent patients infected with natural polio virus (PV), oral PV-vaccinated and inactivated PV-boosted healthy subjects [15– 17], and, most recently, an anti-amyloid antibody with the anti-biofilm activity [18] from a hybridoma library generated with memory B cells from an Alzheimer’s Disease patient [19]. In the current report, eleven hybridoma libraries were generated from the memory B cells of six COVID-19 patients. These libraries were comprised of more than 150 distinct monoclonal antibodies that were selected on the basis of their binding to multiple SARS-CoV-2 proteins in both cell-based and target-based screens. Characterization of these antibodies revealed broad responses to diverse viral antigens. Fewer than half of the antibodies were directed at S protein, while the remainder were directed at other viral proteins including N and ORF-encoded proteins. Even though the antibodies were directed at highly diverse SARS-CoV-2 antigens, they were generally characterized as having variable levels of somatic hypermutation (SHM) and a diversity of V_L_ and V_H_ gene usage. Functional properties of anti-Spike antibodies were successfully confirmed in a pseudovirus neutralization assay. These results indicate that an unbiased interrogation of COVID-19 patient B cell repertoires is an effective approach to identifying specific anti-viral antibodies and antibody mixtures with the desired binding and functional properties. Antibodies identified and characterized in this manner could be recombinantly produced to yield therapeutic or prophylactic products to address the COVID-19 pandemic. The rapidity with which antibodies from convalescent patients can be identified and characterized suggests that this platform could be a useful component of a rapid response to future pandemics.

## RESULTS

### Evaluating the breadth of patients’ humoral responses against SARS-CoV-2

We examined the overall spectrum of the productive antibody response to SARS-CoV-2 using an automated, high-throughput hybridoma library generation and screening platform [15] after isolating memory B cells acquired from blood samples of COVID-19 convalescent patients who demonstrated a high (2880) antibody titer to N and/or S proteins. More than 17,000 hybridomas were generated from the memory B cells of six patients (Table 1), and the naturally occurring human antibodies (IgM, IgG, and IgA isotypes) secreted by those hybridomas were screened for reactivity against a panel of SARS-CoV-2 proteins. Antibody screening assays were developed for three structural proteins (S, N, M) and a panel of accessory ORF proteins of SARS-CoV-2 (Table 2). The screening assays included a rapid and sensitive homogeneous time resolved fluorescence (hTRF) assay that used soluble recombinant viral proteins (Figure 1), as well as a selective, cell-based flow cytometry assay that allowed probing of antibodies to transiently transfected viral antigens expressed within the context of human cells (Figure 2). Commercially available antibodies specific for SARS-CoV-2 proteins demonstrated selective and saturable binding in both assays. The dynamic ranges varied among the specific targets and between assays (Figure 1 and Figure 2) within a 10 pM-100 nM window. Viral protein expression in the cell-based assay was additionally confirmed by Western blot. For cell-based assays, using commercially available controls, the localization of the C-terminus truncated Spike protein (S delta 19aa) was confirmed to be on the surface of the transfected cells, while the localization for Nucleocapsid, Membrane, ORF3a, ORF6, ORF7a, ORF8, and ORF10 proteins was predominantly intracellular.

**Table 1:** Convalescent plasma donors

| Patient ID | Gender | Age |
| --- | --- | --- |
| T_00292 | Male | 31 |
| T_00293 | Female | 44 |
| T_00294 | Female | 34 |
| T_00295 | Female | 53 |
| T_00298 | Female | 46 |
| T_00302 | Male | 45 |

**Table 2:** SARS-CoV-2 viral target panel for screening

| SARS-CoV-2 Viral Targets |  | Assay System |  |
| --- | --- | --- | --- |
| Spike (S) |  | HTRF | Cell-based |
| 1 | Trimer-stabilized FL (non-S1) | ✓ |  |
| 2 | S1 domain (non-RBD) | ✓ |  |
| 3 | Receptor binding domain (RBD) | ✓ | ✓ |
| 4 | S delta 19aa |  | ✓ |
| Nucleocapsid (N) |  | ✓ | ✓ |
| Envelope (E) |  | ✓ | ✓ |
| Membrane (M) |  | ✓ | ✓ |
| ORF3a |  | ✓ | ✓ |
| ORF6 |  |  | ✓ |
| ORF7a |  |  | ✓ |
| ORF8 |  | ✓ | ✓ |
| ORF10 |  |  | ✓ |

**Figure 1.**
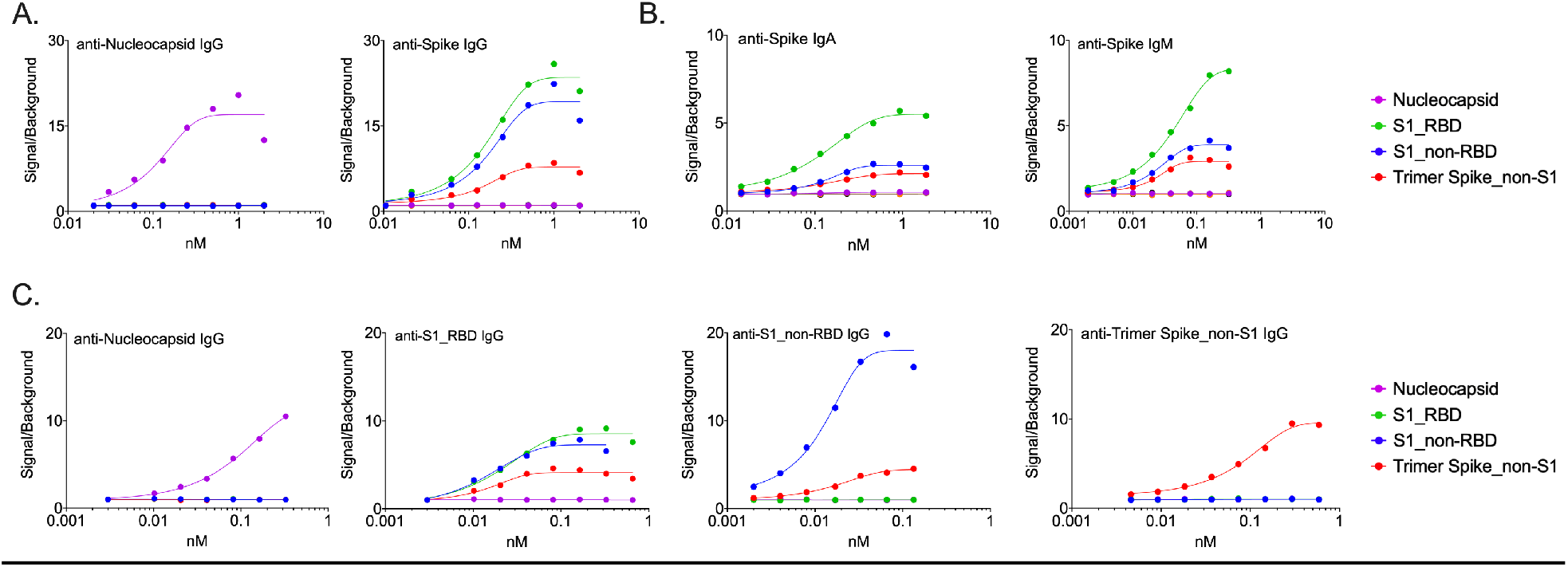
Homogeneous Time Resolved Fluorescence Assay (hTRF) for detection of anti-viral antibodies. (A) Dynamic range and antigen specificity exhibited by Spike and Nucleocapsid control antibodies. (B) Detection of various Spike protein domains with control antibodies of varying isotypes in the hTRF assay. (C) Patient-derived antibodies exhibit specificity and high affinity for target proteins in the hTRF assay.

**Figure 2.**
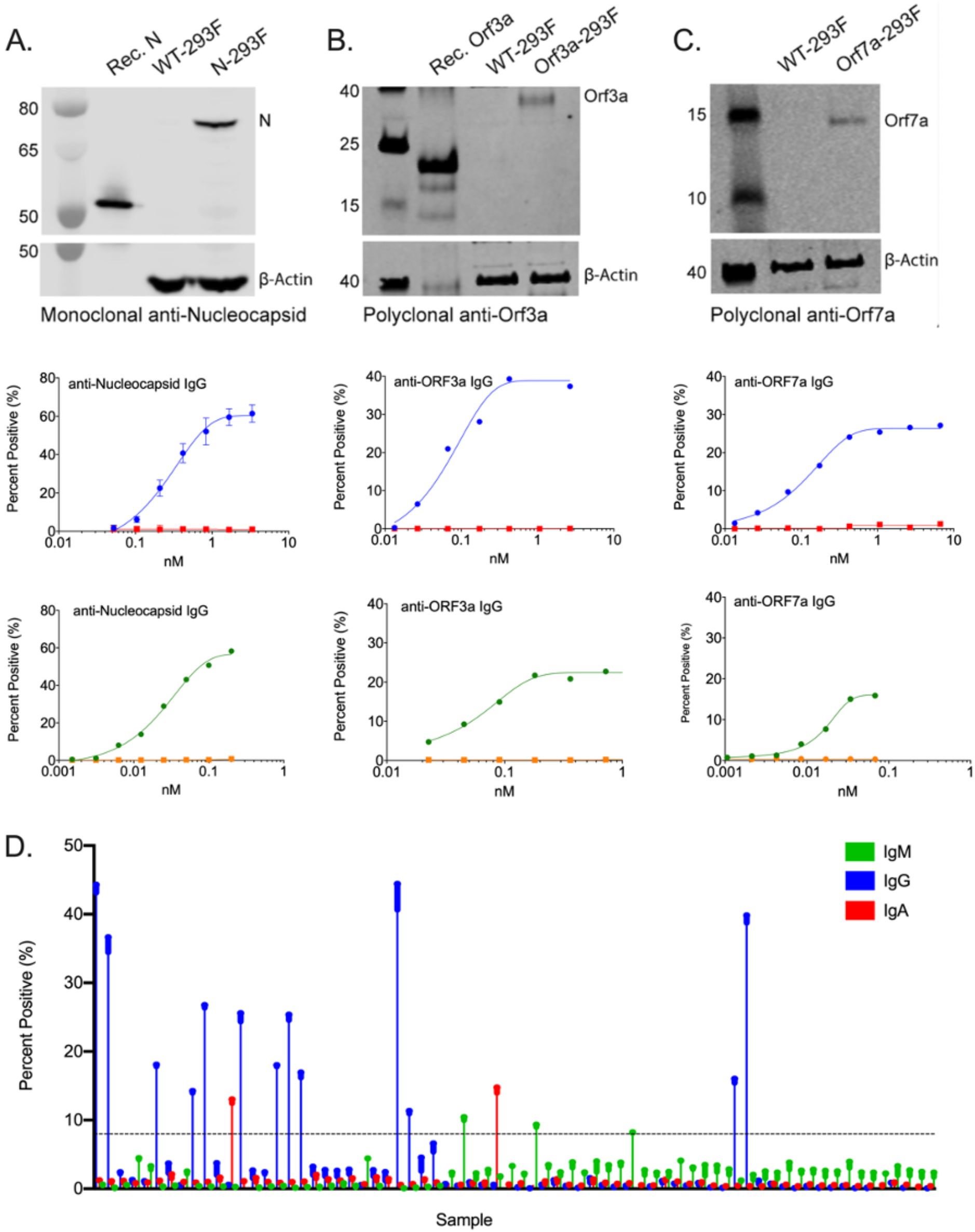
Detection of viral proteins in a cell-based expression system. Commercially available control antibodies were used to detect SARS-CoV-2 protein expression in transiently transfected cells by Western blot (top row) and in a concentration-dependent manner by flow cytometry (middle row). Where available, recombinant protein was used as a positive control for Western blots. Mock-transfected cells served as a negative control for flow-cytometry studies. Patient-derived antibodies demonstrate specificity to antigen-specific cell lines relative to mock-transfected cells (bottom row). (A) Nucleocapsid protein. (B) ORF3a protein. (C) ORF7a protein. (D) Primary screening data depicting isotype-specific detection of patient-derived anti-spike antibodies by flow cytometry.

### Convalescent patients develop antibodies against a broad array of viral proteins

Consistent with previous reports [12,14], S protein was identified as the major antigen for antibody responses in these patients. As shown in Figure 3A, S represented the single most frequently identified target: 76 (44%) of the mono-specific antibodies bound selectively to S protein. As expected, the epitopes for anti-Spike antibodies were distributed across the RBD, non-RBD S1, and non-S1 regions of the Spike protein (Figure 3B). However, consistent with the hypothesis that patients would be mounting antibody responses to a broad range of SARS-CoV-2 proteins, more than half (56%) of identified antibodies bound to non-Spike viral antigens (Figure 3). ORF8 - (28 antibodies, 17%) and N - (23 antibodies, 14%) specific antibodies represented the second and third most-frequently identified targets, respectively. Of note, the antibodies identified in the hTRF assay were biased toward the IgG class (>80%) (Figure 4A). This bias was likely due to lack of sensitivity in the assay for either IgA or IgM, as demonstrated in the control antibody titrations shown in Figure 1. While IgG was still the predominant isotype (∼50%) in the cell-based screening, IgA (∼20%) and IgM (∼30%) antibodies were also well-represented (Figure 4B). Further, these isotypes were also spread across the landscape of antigens being evaluated (Figure 4C).

**Figure 3.**
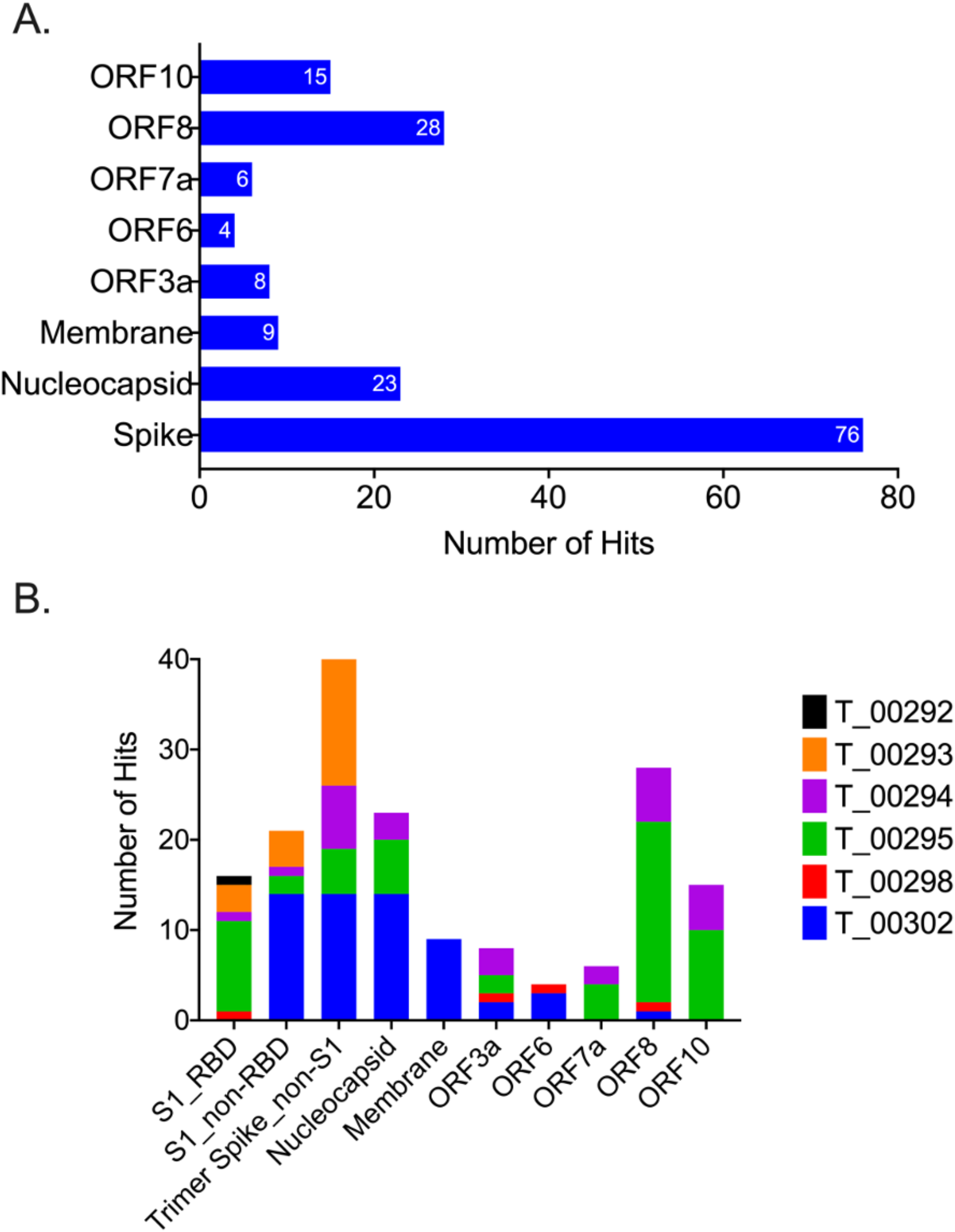
Convalescent patients develop a broad immune response to multiple viral proteins. (A) Less than 50% of the antibodies identified by screening the B cell repertoires of convalescent patients were specific for Spike protein. (B) Individual patients made antibodies against a broad array of SARS-CoV-2 proteins.

**Figure 4.**
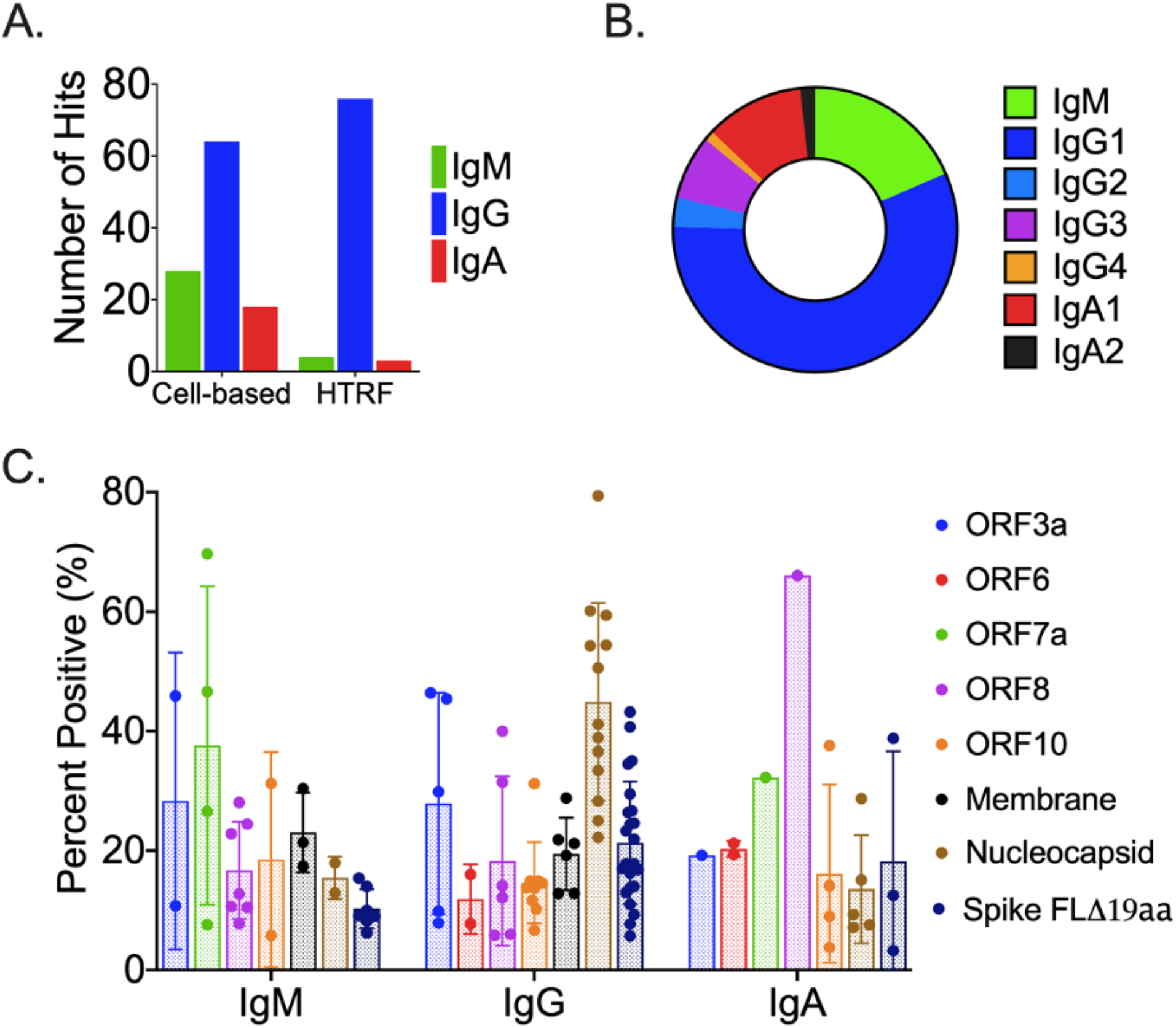
Isotype Distribution of SARS-CoV-2 Specific Antibodies. (A) Distribution identified by the cell- and HTRF-based assays. (B) Distribution of isotypes based upon target. (C) Distribution of isotype sub-classes across targets.

### Immunoglobulin gene usage in convalescent COVID-19 patients

We evaluated Ig gene usage in memory B cells of six COVID-19 patients, using an NGS analysis of identified and sequenced 134 hybridoma hits (Figure 5). Two thirds of identified clones used the kappa locus, while the remaining clones used the lambda locus (Figure 5A). Most of the identified VH regions of IgM, IgG and IgA antibodies belonged to IGHV3 gene family (59%), and the rest were from IGHV4 (15.8%), IGHV1 (14.5%), IGHV5 (7.2%), IGHV2 (2.9%) and IGHV7 (<1%) gene families (Figure 5B). Although there was no dominant V gene variant, IGHV3-21 (8.6%), IGHV3-33 (7.1%) and IGHV3-48 (6.5%) were among the most abundant identified sequences. The most frequent light chain V regions were IGKV3 (26.6%), IGKV1 (22.3%), IGLV3 (13.67%) and IGLV2 (12.2%) (Figure 5C). Furthermore, sequence analysis of hybridoma hits targeting SARS2 S protein (Figure 5D, E) revealed the significant heterogeneity for V gene usage among B cells that produced anti-viral antibodies. This observation is consistent with the parallel expansion of B cell clones targeting multiple components of the SARS-CoV-2 virus, and points to distinct successful strategies employed by the host adaptive immune response during its co-evolution with the virus.

**Figure 5.**
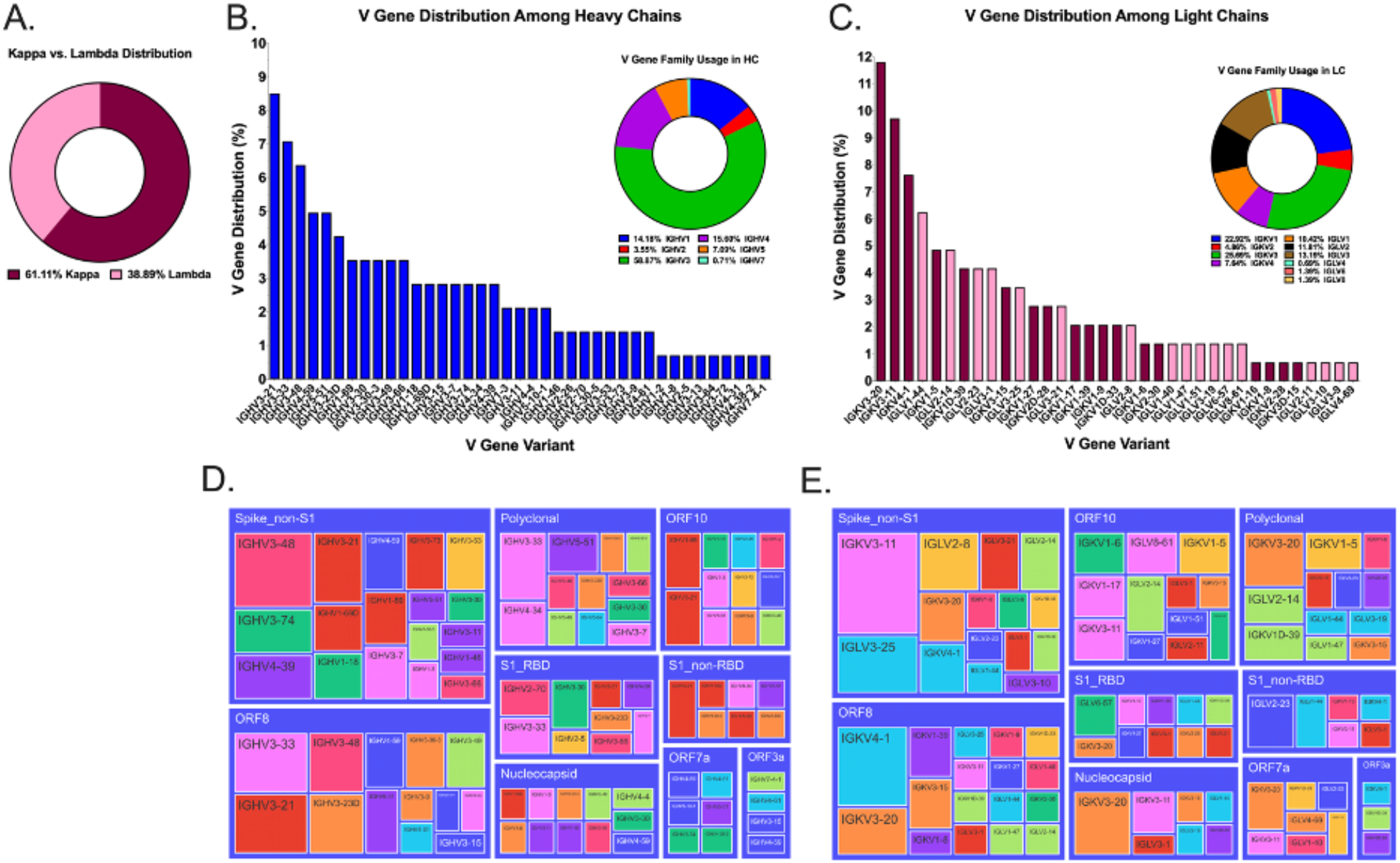
Ig Gene Usage of Identified Screening Hits. (A) Approximately 60% of the identified anti-SARS-CoV-2 screening hits elicited by convalescent patients utilized the kappa light chain locus. (B) VH1 – VH7 gene families were identified as part of antibodies selective for SARS-CoV-2 proteins. (C) Broad range of both kappa and lambda light chains comprise the anti-SARS-CoV-2 primary screening hits isolated by screening the memory B cell repertoires of convalescent patients. (D, E) HC and LC variable domain usage was displayed in a tree diagram using the plotty-express module of the plotty-py program.

### Somatic hypermutations in antibodies from high-titer convalescent COVID-19 patients

We next analyzed heavy and light chain pairs from 103 clones from which we determined productive immunoglobulin RNA sequences. Strikingly, there was a lower-than-expected rate of somatic hypermutation (SHM) in the anti-S antibodies (Figure 6A) and a higher SHM rate in antibodies specific to other viral proteins (ex. N, M, ORF8 and ORF10) (Figure 6A). Of note, even a relatively modest rate of germline mutations in anti-S antibodies resulted in the high affinity Spike-specific antibodies that potently neutralized both S-pseudovirus (Figure 7) and SARS-CoV-2 live virus (manuscript in preparation). Further, a combined analysis of Ig isotype and their level of SHM of virus-specific antibodies revealed several key properties of the productive antiviral response. First, among all “mutated” Igs that had more than 2% of their nucleotide sequence deviated from the closest germline, there was an unusually high (26.4%) proportion of mutated IgMs (Figure 6B), having a mean SHM rate of 5.73%. The functional basis of this phenomenon is not known, but one could speculate that these IgMs came from non-switched memory B cells that had undergone affinity maturation. Second, a subset of such somatically hypermutated IgMs recognized full-length Spike, but not the soluble RBD or S1 subunit of Spike protein. And third, while the predominant isotype among class-switched antibodies was, as expected, IgG (Figure 6C), we were able to capture a panel of fairly mutated virus-specific IgAs. It is plausible that these antibodies play a major role in mucosal neutralization of the incoming virus and may be of particular use for prophylaxis of viral infection and for vaccine design (Figure 6D).

**Figure 6.**
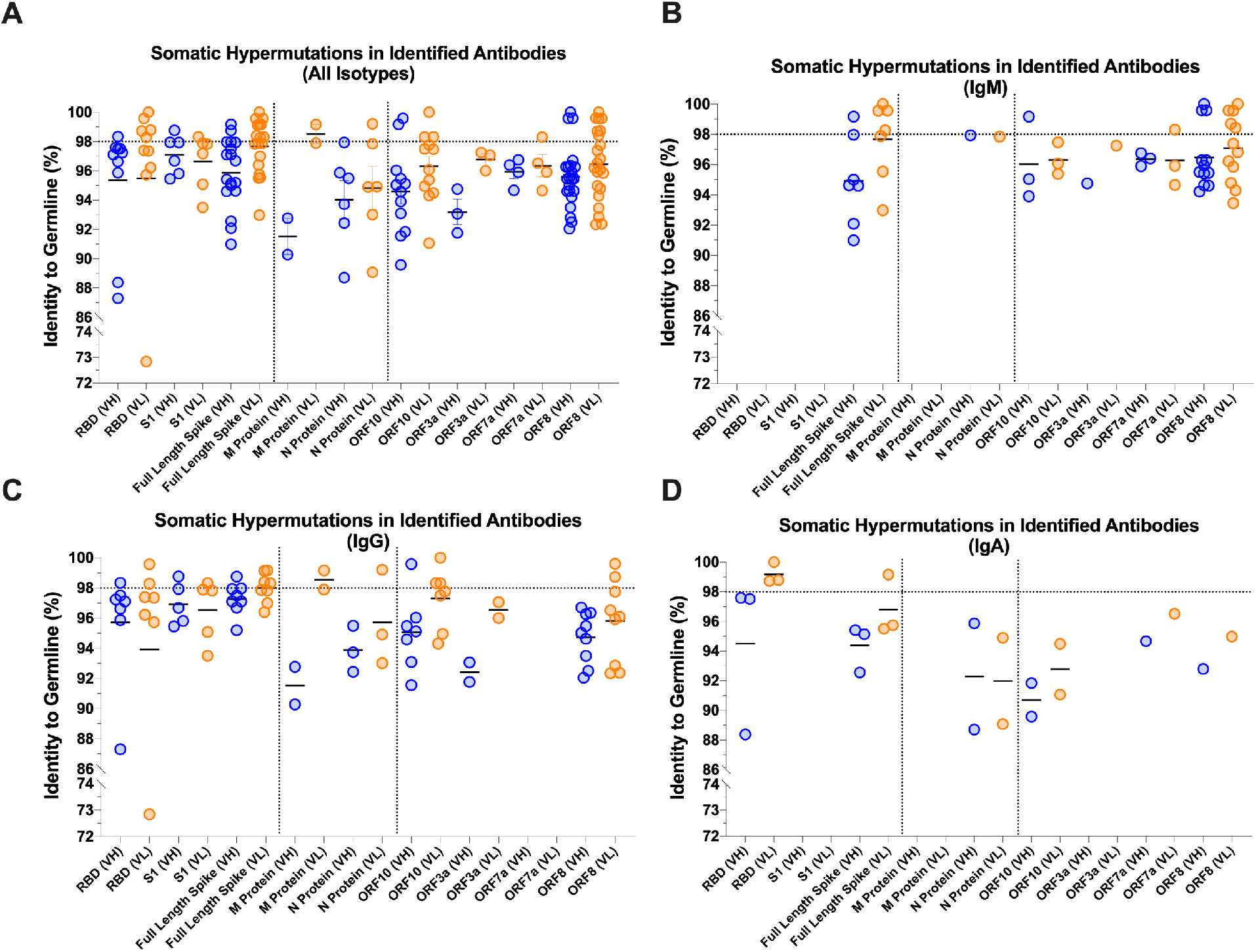
Level of Somatic Hypermutation (SHM) Detected in SARS-CoV-2 Antibodies. (A) Overall and (B-D) isotype-specific levels of SHM in HC/LC pairs associated with the primary screening hits specific for individual viral proteins.

**Figure 7.**
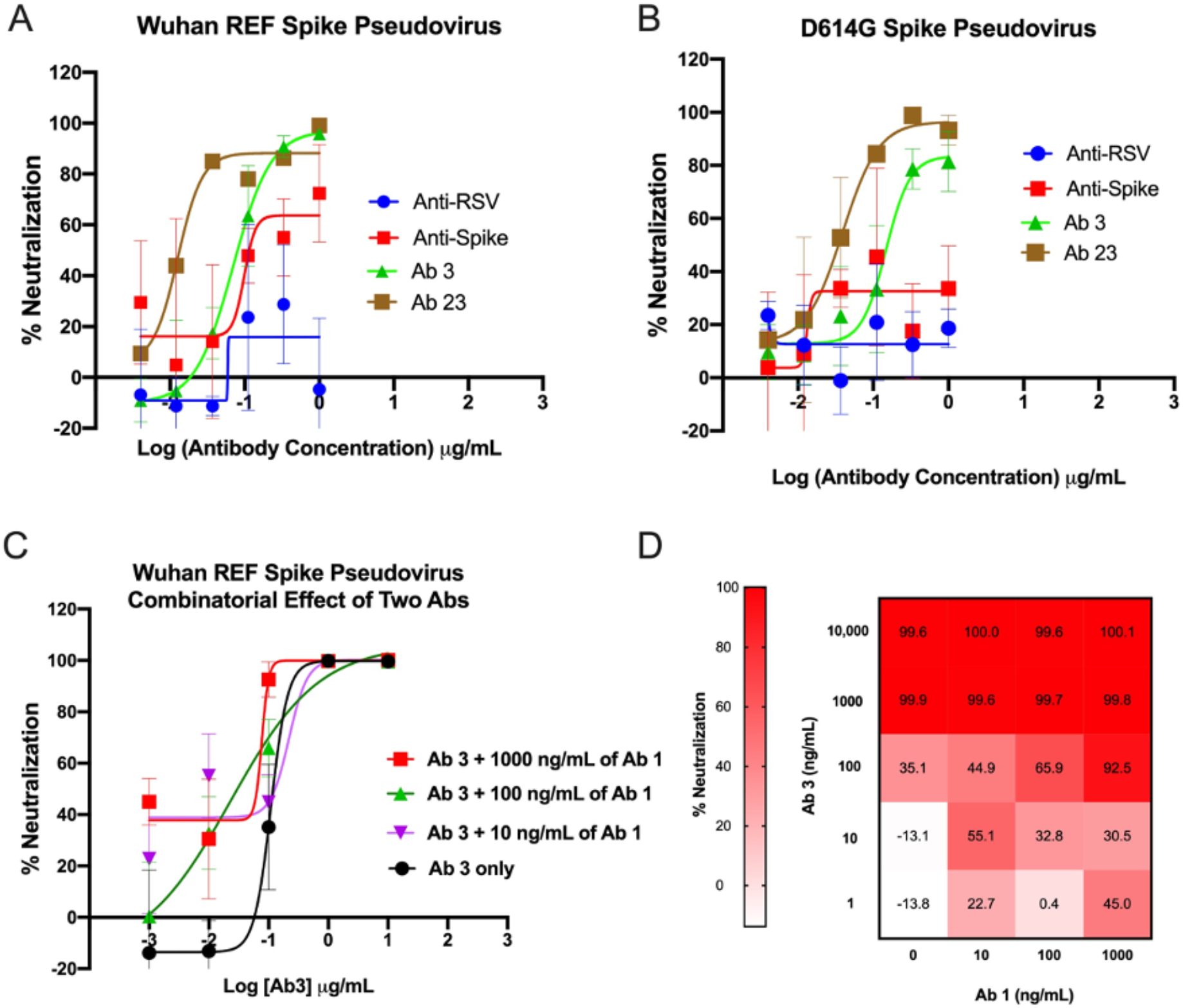
Neutralizing activity of the identified anti-Spike antibodies. (A) Neutralization of WT Spike pseudovirus by selected anti-Spike antibodies. (B) Neutralization of D614G Spike pseudovirus by selected anti-Spike antibodies. (C) Combinatorial effect of Ab#1 and Ab#3 mix on neutralization of WT Spike pseudovirus. (D) Heatmap of percent of neutralization data from panel.

### Identified anti-Spike antibodies from convalescent COVID-19 patients exhibit potent neutralizing activity to Spike pseudovirus

We tested the functional properties of identified anti-Spike antibodies (Figure 7) in a WT and D614G pseudovirus neutralization assay. Spike-expressing pseudoviruses were generated using a lentiviral system and used to infect HEK293 cells overexpressing Angiotensin converting enzyme 2 (ACE2). In the initial screen of antibody-containing supernatants, 3-4 log dilutions of 26 unique anti-Spike antibodies were tested for their ability to block infection in comparison to an anti-RSV negative control and a commercially available anti-Spike positive control. Several antibodies, including Ab#3 and Ab#26, potently neutralized pseudovirus infection with EC50 <500 ng/mL (Table 3). Indeed, a full dose response of purified antibodies confirmed strong neutralizing activity of both antibodies (Figure 7A). In addition to the WT (Wuhan) reference strain, various mutations in the Spike protein of SARS-CoV-2 are prevalent throughout the population, with the D614G variant being the dominant strain at the moment of writing of this manuscript [29]. Thus, identified neutralizing antibodies were tested against and neutralized the D614G S pseudovirus with comparable EC_50_ values (Figure 7B). To further enhance neutralizing activity, combinatorial studies looked for any additive effects seen with the addition of anti-Spike antibodies to Ab#3. Ultimately, Ab#1 at various doses was able to enhance neutralizing activity above that of Ab#3 alone (Figure 7C, D).

**Table 3:** Binding properties of anti-Spike antibodies

| Antibody # | Target<br>(RBD, S1, Trimer S) | *in vitro* EC<sub>50</sub> nM (TRF or cell-based assay) |  | Neutralization potential |
| --- | --- | --- | --- | --- |
|  |  | TRF assay | Cell-based assay |  |
| 3 | RBD, S1, Trimer S | 0.015 | 0.231 | HIGH |
| 26 | RBD, S1, Trimer S | 0.12 | 0.323 | HIGH |
| 10 | RBD, S1, Trimer S | 0.139 | NB | LOW |
| 1 | RBD, S1, Trimer S | 0.021 | 1.11 | MED |
| 2 | RBD, S1, Trimer S | 0.025 | 0.198 | MED |
| 15 | RBD, S1, Trimer S | 0.097 | 1.04 | MED |
| 4 | S1, Trimer S | 0.015 | 0.434 | LOW |
| 5 | S1, Trimer S | 0.111 | 0.198 | LOW |
| 13 | S1, Trimer S | 0.054 | 0.421 | MED |
| 6 | Trimer S | NB | 1.18 | MED |
| Anti-Spike (+) | RBD, S1, Trimer S | 0.904 | ND | 0.316 µg/mL |
| Anti-RSV (-) | N/A | NB |  | 14.125 µg/mL |
|  |  | LOW: neutralization is comparable or below to anti-RSV |  |  |
|  |  | MED: neutralization is observed, but inferior to anti-Spike (Sino Bio) |  |  |
|  |  | HIGH: neutralization is superior to anti-Spike (Sino Bio) |  |  |
|  |  | NB: not binding detected |  |  |
|  |  | ND: not determined |  |  |

## DISCUSSION

The SARS-CoV-2 pandemic has stimulated extraordinary efforts to study anti-viral response and to develop means for treatment and prophylaxis, in both the academic and the biopharmaceutical research communities. By the end of October 2020, a mere 11 months after the virus was first identified, the Clinicaltrials.gov database listed more than 3500 distinct clinical trial activities directed at patients infected with SARS-CoV-2. There is significant diversity among these efforts, from the assessment of existing drugs, to the use of convalescent plasma from recovered patients, to the use of specific vaccines and antibodies directed at the viral S protein. It is not yet apparent that there will be a single approach that will prove to be uniformly effective at preventing viral infections or accelerating viral clearance in all groups of COVID-19 patients.

It is possible that even the most effective approaches will have limited or unsustainable efficacy. The vast majority of the ongoing efforts are all targeting the S protein. Both passive (therapeutic antibodies) and active (vaccine) approaches directed at S protein are expected to promote virus neutralization, that is, inhibition of viral entry into healthy cells. Unfortunately, a mutation in S protein has already been reported [20,21] and further mutations may ultimately limit the effectiveness of therapies directed at this single protein [22].

Given the multiplicity of SARS-CoV-2 proteins that induce antigen-specific T and B cell responses in humans, it is reasonable that an unbiased interrogation of the memory B cells generated by high-titer, convalescent COVID-19 patients could identify high-affinity Igs directed at specific viral antigens. As an outcome of this approach, there was not a single dominant V gene LC/HC combination specific to a particular viral protein among characterized 134 Igs from convalescent patients. In fact, sequence analysis revealed a lower rate of SHM of anti-Spike Igs than of antibodies directed at other viral proteins (N, M, ORF8 and ORF10). Remarkably, even a modest rate of germline mutations in anti-S antibodies resulted in potent neutralization of both S-expressing pseudovirus (Figure 7) and SARS-CoV-2 live virus (manuscript in preparation). To our surprise, these Spike-specific antibodies included an atypically high proportion of well-mutated anti-viral IgMs (26.4% with a mean mutation rate of 5.73%) suggesting that non-switched memory B cells also undergo affinity maturation in response to SARS-CoV-2 infection. Further, a subset of such mutated IgMs targeted the full-length Spike, but not soluble S1 or RBD domains. The identification of IgG and IgA antibodies specific for the full range of targets screened, including the RBD domain of S, highlights the requirement of additional events, such as immunoglobulin class switching, for the development of a productive neutralizing antibody response. Finally, and perhaps not unexpectedly, we have identified a group of polyreactive antibodies (data not shown). A relatively high rate of SHM in these polyreactive antibodies may suggest a secondary maturation event that redirected immature B cell clones toward SARS-CoV-2 antigens.

The majority of anti-S antibodies, as expected, recognized a full-length S protein (Table 3). Further, all but one of the six COVID-19 convalescent patients produced antibodies to a soluble receptor-binding domain of S protein (RBD). Of note, while most of the RBD- and S1-specific antibodies were less mutated than those specific to the full-length S protein, we identified several highly mutated RBD-specific outliers. Lower rate of SHM in the S-specific antibody group may connote limited rounds of affinity-maturation for a high antigenic protein. These anti-Spike antibodies demonstrated functional activity in a pseudovirus neutralization assay. In fact, there were several potent neutralizing antibodies (e.g., Ab#3, Ab#26), that had EC_50_ in 100 ng/mL range. One of these two antibodies demonstrated a combinatorial effect with Ab#1 in pseudovirus (Figure 7C, D) neutralization assays. The selectivity, affinity, and functional activity of the anti-Spike antibodies suggest that they were a part of the successful anti-viral responses mounted by the patients from which they were derived. By extension, the antibodies identified against other viral targets may have also contributed to the viral clearance through mechanisms other than neutralization (i.e., activators of complement and/or effector cells).

In summary, an unbiased interrogation of the B cell repertoires of convalescent COVID-19 patients demonstrated that these patients make a strong humoral response against a broad array of SARS-CoV-2 proteins. These responses included high affinity antibodies of multiple Ig isotypes. The natural immune response to SARS-CoV-2 among these patients stands in stark contrast to the anti-S focused approaches being taken to develop therapeutic antibodies to treat COVID-19. An alternative approach should include targeting a breadth of SARS-CoV-2 proteins with a cocktail of antibodies, with the anticipation that a multi-targeted mixture will be more effective at inducing robust viral clearance via neutralization and Fc-mediated activation of complement and effector cells.

## MATERIAL AND METHODS

### Cells

293TN Producer cell line (System Biosciences, Cat #LV900A-1) was maintained in DMEM containing 10% FBS. HEK293 cells expressing human Angiotensin converting enzyme 2 (ACE2) (BPS Biosciences, Cat #79951) were cultured in EMEM containing 10% FBS and 5 mg/mL Puromycin to select for ACE2-expressing cells. ACE2 expression was confirmed by flow cytometry. Cells were detached with CellStripper (Corning, Cat # 25-056-Cl) and labeled with LIVE/DEAD Aqua (Invitrogen, Cat # L34966) at 1:1000 dilution at room temperature for 10 minutes. After that, cells were washed twice in PBS and stained with a goat anti-human ACE2 polyclonal antibody (R&D Systems, Cat # AF933) or isotype goat polyclonal isotype control (R&D Systems, Cat # AB-108-C) at 1:100 dilution for 30 minutes on ice. Next, cells were washed twice and stained with Alexa Fluor 488-conjugated donkey anti-goat antibody (Jackson ImmunoResearch, Cat #705-545-147) at 1:200 dilution for 30 minutes on ice. Finally, cells were washed twice and run on the Attune NxT (ThermoFisher). Data were analyzed using FlowJo software (BD).

### Collection of patient samples

Blood samples were drawn from six convalescing COVID-19 patient volunteers deemed eligible for donating convalescent plasma as set forth in the US FDA’s Recommendations [23]. Patients displayed no PCR-detectable viremia and maximal IgG (2880) titer of class-switched, virus-specific antibodies. Donors gave written consent to have their blood drawn and authorized the unrestricted use of their blood samples by Immunome. The samples were deidentified and the B cells were isolated from those deidentified blood samples. Immunome did not seek IRB approval of a “research project” because analysis of peripheral blood samples that are obtained with consent, deidentified, coded or anonymized are not believed to be subject to human tissue research regulations. Immunome used a commercial vendor to obtain additional blood samples, one with a standing IRB approval in place for their donor collection efforts.

### Generation of hybridoma libraries

Hybridomas were generated following protocols for isolating and expanding primary B-cells as well as electrofusion methods described in U.S. patents [24,25]. Hybridomas stably expressing human mAbs were generated by electrofusion of expanded B-cells to the B5-6T myeloma cell line, which expresses an ectopic human telomerase gene that stabilizes human chromosomes in the hybrid cells created. Fused hybridomas were plated into 96-well plates in growth medium with HAT selection of stable hybridomas for 7 days. After 7 days, growth media were switched to media with HT for stable selected hybridoma growth. Hybridomas were cultured in a 37°C incubator for 14-21 days during which time they were imaged for monoclonality and monitored for isotype-and sub-class-specific Ig secretion. Supernatants from monoclonal wells expressing measurable levels of Ig were cherry-picked and submitted for target-based screening.

### HTRF Screening Assays

A homogeneous time-resolved fluorescence (hTRF) assay [26] comprised of terbium-labeled anti-human IgG (H+L) (Cisbio, custom label) donor and AF488-labeled anti-HIS (Cell Signaling, Cat # 14930S) acceptor antibodies was used to screen patient-derived antibodies for their binding to recombinantly produced SARS-CoV-2 antigens. The assay, adapted for high-throughput screening, was optimized so that a number of recombinant, HIS-tagged SARS-CoV-2 target proteins could be substituted interchangeably. This recombinant target panel consisted of two full-length viral structural proteins, S (FL, trimer-stabilized, LakePharma) and N (GenScript); two truncated S protein domains, S1 (GenScript, Cat # Z03485-1) and RBD (aa 319-591, LakePharma); and two ORF proteins, ORF3a (ProSci, Cat # 10-005) and ORF8 (ProSci, Cat # 10-002). Commercially available antibodies specific for the individual structural viral proteins, SARS-CoV/SARS-CoV-2 Spike S1 (RBD) chimeric mAb (Sino Biological, Cat # 40150-D001), SARS CoV-2 Nucleocapsid human chimeric mAb (GenScript, Cat # A02039-100), or in-house antibodies to ORF3a and ORF8 served as positive controls. Assay background was determined by averaging the signal of wells containing only the donor and acceptor cocktail. Hybridoma supernatants exhibiting signals greater than 2-fold over background were reported as positive HITs and are submitted for Ig sequence analysis.

### Flow cytometry-based cellular screens for antiviral antibodies

SARS-CoV-2 antigen sequences were cloned into pcDNA3.4 plasmids and transfected into 293F cells utilizing the Expi293 Expression System (Life Technologies, Cat # A14635) per manufacturer’s instructions. Cells expressing SARS-CoV-2 N, ORF6, ORF8, and ORF10 proteins were transferred into Expi293 Expression medium (Life Technologies, Cat # A1435103) under antibiotic (Geneticin) selection (Gibco, Cat # 10131027) for 3-5 days following transfection to establish individual stable cell pools. The stable cells were maintained under selection in the presence of Geneticin. Protein localization was confirmed by flow cytometry for either intact or fixed and permeabilized Intracellular Perm Buffer (BioLegend, Cat # 421002)) cells. Cells transiently expressing SARS-CoV-2 S (S delta 19aa), M, ORF3a, and ORF7a were included in the screening panel. Optimal protein expression was achieved two days post-transfection for ORF3a and ORF7a and three days post-transfection for S and M proteins.

For both the cell-surface binding and intracellular detection assays, cells were incubated with LIVE/DEAD Cell Stain kit (ThermoFisher, Cat # L34960) per the manufacturer’s instructions. For cell surface binding, live cells expressing S delta 19aa were suspended in QSol Buffer (IntelliCyte Corporation) to which a cocktail of Fc-specific secondary antibodies was added: AF647 goat anti-human IgG (Jackson ImmunoResearch, Cat # 109-605-008), AF488 goat anti-human IgA (Jackson ImmunoResearch, Cat # 109-115-011) and BV650 mouse anti-human IgM (BioLegend, Cat # 314526).

For the permeabilization assay, stained cells were first fixed with paraformaldehyde (BioLegend, Cat # 420801) at a final concentration of 1%. Fixed cells were then permeabilized with Intracellular Permeabilization Buffer (BioLegend) according to the manufacturer’s instructions. A cocktail of Fc-specific secondary antibodies consisting of AF647 goat-anti-human IgG (Jackson ImmunoResearch, Inc.), PE goat-anti-human IgA (Jackson ImmunoResearch, Inc.), and BV650 mouse-anti-human IgM (BioLegend) was added to cell suspension.

For each assay, the cell suspensions were dispensed into 384-well plates, followed by the addition of hybridoma supernatant at a 1:10 final dilution. The reaction was allowed to incubate for 90 minutes at room temperature. For the permeabilization assay, plates were centrifuged, and cell pellets were suspended in QSol Buffer (IntelliCyte Corporation). Cells for each assay were fixed with a final concentration of 1% paraformaldehyde and analyzed with IntelliCyte iQue Screener (IntelliCyte Corporation). Positive binding gates for detection of each secondary antibody were established using cells plus secondary antibody cocktail as a negative control. Binding of hybridoma supernatant antibodies to specific SARS-CoV-2 proteins was quantified as percent positive relative to the secondary only control. To calculate percent positive, live events that shifted into detection channels were divided by the total live events.

### RNA isolation and Next Generation Sequencing (NGS)

Hybridoma RNA was isolated using RNAqueous-96 Total RNA Isolation Kit (Invitrogen, Cat # AM1920). Isolated RNA samples were submitted to iRepertoire (Huntsville, AL) for NGS. Hybridoma-derived RNA samples were sequenced using the Illumina MiSeq system at iRepertoire (Huntsville, AL). Sequencing runs were performed using the MiSeq Nano Kit V2 following bead-based cleanup of RNA. Immunoglobulin sequences containing CDR1, 2 and 3 and framework regions were amplified using IgG and IgA-specific mixes for IgH, and kappa and lambda-specific primers for IgL. IgM-expressing hybridoma samples, from which IgG or IgA heavy chains were not amplified using this approach, were sequenced using the iRepertoire iPair system. Final sequences were exported using iPair software.

The analysis of primary NGS data was performed by iRepertoire. Immunoglobulin sequences were analyzed for predicted CDR sequences, % identity to appropriate germlines, isotype of the constant regions and read counts. Sequence pairing was performed based on the read count information. In the event more than one LC:HC pair was discovered in a single well, each LC:HC combination was analyzed as a separate antibody. In wells exhibiting 5’ truncation in the V region, the germline sequence was used to create an expression construct. Final sequences were translated and analyzed for potential stop codons and frame shifts.

### Production of paired light and heavy chains

Variable domains yielding productive uninterrupted protein sequences were analyzed for number of reads and the degree of somatic hypermutations (SHM) in comparison to the closest immunoglobulin germline. Hybridoma hit sequences with at least one chain that had more than 2% of SHM were advanced to HC/LC pairing and the recombinant production of antibodies. Immunoglobulin expression fragments were cloned into the pcDNA3.4-based vectors and expressed in 293F cells. Affinity and binding pattern of recombinant antibodies were compared to the original antibody-containing hybridoma supernatants in BLI, HTRF and cell-based assay. Antibody-containing supernatants or purified antibodies were advanced to downstream assays. If multiple heavy or light chain sequences were detected within one well, their CDRs were aligned and compared for potential PCR errors. In cases where multiple sequences within a well were different, i.e., originated from separate clones, all potential combinations of light and heavy chains were recombinantly produced and tested in downstream assays. Wells that yielded a single HC/LC pair were advanced to recombinant expression and downstream assays. 5’ fragments of the constant regions were sequenced to identify the isotype of the antibody and compared to the experimentally identified isotype of hybridoma supernatants. The resulting isotype of the heavy or light chain was assigned based on two or more positive readings from experimental (ex. ELISA and FACS) assays and sequencing.

### Pseudovirus Production and Neutralization Assay

Spike-expressing pseudovirus was generated with System Bioscience’s pPACK-SPIKE packaging system (System Biosciences, Cat # CVD19-50A-1) as per manufacturer’s protocol. Briefly, 8×10^6^ 293TN Producer cells (System Biosciences, Cat # LV900A-1) were plated in T150 flasks overnight. Plasmids encoding lentiviral packaging proteins and Spike were added 1mL of plain DMEM for each T150 being transfected. 55 mL of PureFection reagent (System Biosciences; Cat # LV750A-1) was added to each 1mL tube, vortexed for 10 seconds, and incubated at room temperature for 15 minutes. The plasmid and PureFection mixture was added to a T150 flask containing 293TN cells and placed in a 37°C incubator containing 5% CO_2_ for 48 hours. Pseudovirus-containing supernatants were harvested at 48 hours and passed through a 0.45-micron PVDF filter to remove cellular debris. 5x PEG-it Virus Precipitation Solution (System Biosciences, Cat # LV810A-1) was added to supernatants and incubated 4°C overnight. Pseudovirus-containing supernatants containing 1x PEG-it Virus Precipitation Solution were then spun at 1500 x g for 30 minutes. Pseudovirus-containing pellets were resuspended in plain DMEM to achieve at 10x concentration and frozen at -80°C in single use aliquots.

Pseudovirus infection and neutralization assays were performed by adapting established protocols [27, 28]. In summary, 10^4^ ACE2-293T cells were plated in the inner 60 wells of a 96 well flat bottom plate in 100uL of ACE2-293T media overnight in a 37°C incubator containing 5% CO_2_. To determine infectivity of each lot of pseudovirus, pseudovirus-containing supernatants were thawed from -80°C and two-fold dilutions were performed. 100 mL of pseudovirus at various dilutions was added to ACE2-293T cells. To test neutralization activity of antibodies, indicated antibody concentrations were pre-incubated with pseudovirus for 1 hour in a 37°C incubator containing 5% CO_2_. Then, 100 mL of antibody/pseudovirus mixture was added to ACE2-293T cells. After 72 hours, cells and media were equilibrated to room temperature for 20 minutes. 100 mL of media was removed and replaced with 100 mL of Bright-Glo Luciferase Assay Reagent (Promega, Cat # E2620). Luciferase activity was measured on the EnSpire Plate Reader (PerkinElmer). Percent neutralization was calculated using RLUs with the equation [(RLU of Virus + cells) – (RLU of Experimental Sample)] / [(RLU of Virus + cells) – (RLU of cells only)].

## ACKNOWLEDGEMENTS

This study was funded by the U.S. Department of Defense (DOD) Joint Program Executive Office for Chemical, Biological, Radiological and Nuclear Defense (JPEO-CBRAND) Joint Project Manager for Chemical, Biological, Radiological and Nuclear Medical (JPEO-CBRN Medical), in collaboration with the Defense Health Agency (DHA), under contract W911QY2090019. The opinions, interpretations, conclusions and recommendations are those of the authors and are not necessarily endorsed by the U.S. Army.

